# Natural variation in salt-induced root growth phases and their contribution to root architecture plasticity

**DOI:** 10.1101/2023.01.27.525841

**Authors:** Eva van Zelm, Silvia Bugallo-Alfageme, Pariya Behrouzi, A. Jessica Meyer, Christa Testerink, Charlotte M.M. Gommers

## Abstract

During salt stress, the root system architecture of a plant is important for survival. Different accessions of *Arabidopsis thaliana* have adopted different strategies in remodeling their root architecture during salt stress. Salt induces a multiphase growth response in roots, consisting of a stop phase, quiescent phase, recovery phase and eventually a new level of homeostasis. We explored natural variation in the length of and growth rate during these phases in both main and lateral roots and find that some accessions lack the quiescent phase. Using mathematical models and correlation-based network, allowed us to correlate dynamic traits to overall root architecture and discover that both the main root growth rate during homeostasis and lateral root appearance are the strongest determinants of overall root architecture. In addition, this approach revealed a trade-off between investing in main or lateral root length during salt stress. By studying natural variation in high-resolution temporal root growth using mathematical modeling, we gained new insights in the interactions between dynamic root growth traits and we identified key traits that modulate overall root architecture during salt stress.

**Summary statement:** By studying natural variation in salt-induced root growth phases in *Arabidopsis*, we show that main root growth rate during homeostasis and lateral root appearance contribute most to root architecture and we reveal a trade-off between investing in main and lateral root growth during salt stress.

## Introduction

Soil salinization is a worldwide problem, affecting 16.5 MHa of agricultural land between 1980 and 2018 (Hassani et al., 2020), and due to climate change is predicted to increase in many regions in the coming 80 years (Hassani et al., 2021). Soil salinity reduces crop yield and thereby poses a problem for agriculture. In plants, salt causes osmotic stress and ion toxicity, which can limit growth by reducing water uptake and interfering with photosynthetic productivity, respectively [reviewed in (Munns and Tester, 2008; Julkowska and Testerink, 2015; Van Zelm et al., 2020)]. Naturally occurring accessions of the model plant *Arabidopsis thaliana* have been used to characterize variation in both shoot and root growth responses to salt (Julkowska et al., 2014; Awlia et al., 2016). Furthermore, natural variation in *Arabidopsis* accessions has been successfully used to identify candidate genes that affect various salt related phenotypes such as root architecture remodeling (Julkowska et al., 2017), shoot growth (Awlia et al., 2021), sodium accumulation (Rus et al., 2006) and salt avoidance by the main root (Deolu-Ajayi et al., 2019; Zou et al., 2022a), using genome-wide association mapping. These approaches have revealed new pathways that modulate plant responses to salinity and that could eventually influence salt tolerance [reviewed in (Zou et al., 2022b)].

The root is the first organ that encounters salt in the soil and has an important role in both uptake and exclusion of sodium ions. In response to salt the root system architecture is not only retarded in its development, but also remodeled, as the distribution of root mass between main and lateral roots is altered (Julkowska et al., 2014). Studies to natural variation in Arabidopsis accessions suggested that during salt stress a trade-off occurs between investment in either main or lateral roots. In control conditions, main and lateral root growth are positively correlated, but during high levels of salt stress this correlation is lost and instead becomes a negative trend (Julkowska et al., 2014). Furthermore, different accessions of Arabidopsis employ different strategies in root architecture remodeling, resulting in differences in main root length, average lateral root length and lateral root number (Julkowska et al., 2014). This natural variation is mainly due to differences in lateral root emergence and elongation, rather than differences in the patterning of lateral roots (Julkowska et al., 2017). Surprisingly, little is known about the differences in growth rates between main and emerged lateral roots in general or in response to changes in the environment (Waidmann et al., 2020). Here, we aim to investigate whether differences in growth rates of main and emerged lateral roots cause root architecture remodeling during salt stress.

When measured at a high temporal resolution, salt induces interesting growth dynamics in both main and lateral roots in Col-0 (Duan et al., 2013; Geng et al., 2013). In response to salt, the growth rate of the main root is rapidly reduced and enters a phase of minimal growth (quiescent phase) (Geng et al., 2013). After this quiescent phase, the root growth partially recovers and reaches a new homeostasis. These growth phases are regulated by different hormones, such as abscisic acid (ABA), jasmonic acid and brassinosteroids and the crosstalk between them (Geng et al., 2013). Lateral roots enter a prolonged phase of growth quiescence after their appearance in response to salt, a process which is regulated by ABA (Duan et al., 2013). The impact of these growth phases of main and lateral roots on root architecture remodeling is still unknown. We use phenotypic variation amongst accessions of Arabidopsis to understand which growth phases are important for the remodeling of over-all root architecture in salt.

In this work, we study natural variation in salt-induced root growth phases and in the existence of a salt-induced quiescent growth phase for both main and lateral roots amongst accessions of Arabidopsis. Fitting a double sigmoidal equation, allows us to quantitatively distinguish different root growth phases. We show that the dynamics of the recovery and the homeostasis phase correlate between main and lateral roots, while the length of the quiescent phase does not. We correlate the dynamic changes in root growth to overall root architecture, and show that the length of the main root recovery phase, the main root growth rate during homeostasis and the delay in lateral root appearance contribute to overall root architecture during salt stress. Interestingly, the impact of the main root growth rate during homeostasis and the delay in lateral root appearance reveal a trade-off between investing in main and lateral root tissue during salt stress, which confirms the previously suggested trade-off under salt stress (Julkowska et al., 2014).

## Methods

### Plant materials and growth conditions

Based on previously published root architecture traits (Julkowska et al., 2014), accessions with a moderate (falling in the interquartile range) main and average lateral root length in control conditions were selected (supplemental figure 1A). From these, a subset of 20 accessions with either small or large salt-induced changes in main or average lateral root length were selected (supplemental figure 1B). Seeds were surface sterilized in 75 % EtOH and 25 % bleach, after which they were washed twice with 70% EtOH and once with 96 % EtOH. The EtOH was removed and 0.1 % agar was used to sow the seeds on plates containing 0.5 x Murashige-Skoog (MS, Duchefa), 0.1 % M.E.S. Monohydrate (Duchefa) and 1 % Daishin agar (Duchefa) (pH 5.8, with KOH). Plates were left for 7 days at 4 °C for seed stratification and grown under long day conditions (16 h light : 8 h dark, 130 μmol/m^2^/s LED light) at 21 °C and 70% relative humidity.

For measurements of root system architecture traits, seedlings were grown for 5 days at a 70° angle, transferred to plates with or without 125 mM NaCl and scanned 7 days after transfer. Root architectures were traced using SmartRoot 2.0 (Lobet et al., 2011) and the calculated traits were compared to published measurements using a Pearson’s correlation test. Outliers were removed using the rstatix package in R (https://cran.r-project.org/web/packages/rstatix/index.html). For main and lateral root dynamics, seedlings were transferred to control or salt media at 7 or 9 days-old and imaged every 20 minutes with an automated infrared imaging setup, in which plates were positioned at a 90° angle. The main or lateral root length in the last image was traced using SmartRoot and the root length in previous pictures was corrected using an automated edge-detection script [as described in (Gigli-Bisceglia et al., 2022)]. The root length was averaged per hour and the growth rate was calculated over 2 hour intervals. For the turning assay, 5 day-old plants were transferred to plates with or without 125 mM NaCl and turned 90°. Plates were scanned daily and the moment of lateral root appearance was scored manually. Throughout the experiment plates were placed at a 90° angle. For all assays, seedlings that stopped growing after transfer, grew away from the plate or could not be traced accurately were removed from the analysis.

### Fitting growth rate dynamics

A sigmoid is a commonly used model to describe the increase/decrease from a floor to a ceiling value, where a double-sigmoidal model can be used to describe two growth and/or decay phases (Caglar et al., 2018; Baione et al., 2020). Under salt stress, main and lateral root growth rate first decreases and subsequently increases, therefore we used a double sigmoidal model to describe this growth behavior. However, because not all accessions empirically show a decrease phase (stop phase) especially for the lateral root data, we fitted both a single and a double sigmoid and evaluated which model described the data better. Thus, we fitted both a single (1) and a double sigmoidal (2) model to the average growth rate per hour with the following equations:

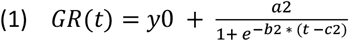

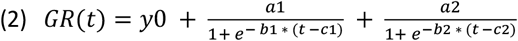

Where GR is the growth rate, y0 is the lower asymptote, a1 and a2 are the decrease in the first sigmoid and the increase in the second sigmoid respectively, b1 and b2 are the slopes of the decay and growth phase of the first and second sigmoid and c1 and c2 are the midpoints of the first and second sigmoid and t is the time after treatment (Figure 2A). Fittings were performed using nlsLM from the minipack package in R (https://cran.r-project.org/web/packages/minpack.lm/minpack.lm.pdf). The b1 did not show significant estimates for any of the accessions. To avoid overparameterization, b1 was fixed to the median of the initially fitted values (b1 = -2.05 for main root, b1 = -0.43 for lateral root). Parameter estimates were not allowed to be lower than 0 and are shown in supplemental figure 5 and 8 for the main and lateral roots respectively. We used the Akaike information criterion (AIC) to determine which model was a better fit for the growth data. The double sigmoid was considered a better fit if AIC for single sigmoid – AIC for double sigmoid > 2.

The local maxima and minima in the second derivative (peaks) of these models were used to define the begin and end point of different growth phases (Figure 2). For the double sigmoid, this resulted in two distinct growth patterns amongst accessions. Accessions showed either 2 or 4 peaks after the midpoint of the first sigmoid (Figure 2A & B). For accessions with 4 peaks after the first midpoint, the 1^st^ peak marked the start of the quiescent phase and the 3^th^ and 4^th^ peak marked the start and the end of the recovery phase (Figure 2C). For accessions with 2 peaks after c1, the peaks were used as indication for the start and the end of the recovery phase (Figure 2C). For growth data where the single sigmoid was a better fit, the 1^st^ and 2^nd^ peak in the second derivative of the sigmoid were defined as the start and the end of the recovery phase and the time before the 1^st^ peak was defined as the quiescent phase.

### Partial correlation network

For all 16 traits, a ratio between salt and control was calculated, except for the durations of the different growth phases. For the growth rate within a growth phase, the average growth rate during salt stress in the timeframe of the phase was divided by the average growth rate in control conditions in the same timeframe. For the accessions that did not show a specific phase, the length and rate of this phase were not taken along in the analysis.

To find simultaneous correlations among all 16 traits, we used mixed graphical models (Behrouzi and Wit, 2019). This model allows us to discriminate between direct and induced correlations which is a key contributor to minimize false positive interactions and leads to a sparse structure. In other words, this model encodes conditional independence (i.e. direct) relationships between the traits, and can be visualized by a network where vertices represent the traits and the conditional dependence among them is visualized by edges. The strength of interactions between traits are determined using the inverse of variance-covariance matrix, where the thickness of edges shows partial correlation values. The method can be used to estimate high-dimensional interactions networks for non-Gaussian, mixed discrete-and-continuous data. The employed methodology is implemented in a publicly available R-package netgwas (CRAN - Package netgwas (r-project.org)) (Behrouzi and Wit, 2019). This exploratory statistical methodology enables to reveal previously unnoticed patterns in datasets and can pinpoint novel mechanisms.

## Results

### Not all Arabidopsis accessions show a main root quiescence growth phase in response to salt

We reasoned that accessions that display different root architecture remodeling under salt stress could be a useful tool to study the relationship between root growth dynamics and overall changes in root architecture under salt. For a subset of accessions with diverse salt-induced root architecture remodeling, we selected 20 accessions based on the differences in main and average lateral root length from a previous study (Julkowska et al., 2017) (supplemental figure 1, explained in the material and method section). Col-0 did not fit our criteria but was included because it is the most used accession in other studies. To reproduce the overall root architecture of these accessions in our experimental setup, we transferred 5-day old seedlings to 125 mM salt or control plates and quantified root architecture traits 7 days later in 12-day old plants. In our experimental setup, this selection of accessions displayed a range of altered main root length, average lateral root length and lateral root number in response to salt stress (Figure 1, supplemental figure 2). To see how robust the salt-induced changes in root architecture are between experimental setups, we compared the changes in main and lateral root length in response to salt between our experiment and the published data (Julkowska et al., 2017) (supplemental figure 3). This analysis indeed showed that the changes in main root length under salt stress correlate between separate experiments amongst accessions (Pearson correlation, R^2^ = 0.54, p < 0.05, supplemental figure 3A). The average lateral root length in response to salt on the other hand does not significantly correlate between experiments (Pearson correlation, R^2^ = 0.38, p > 0.05, supplemental figure 3B). Tsu-0 is an outlier in both correlations and in a different study Tsu-0 showed a root architecture comparable to our measurements (Julkowska et al., 2014). When Tsu-0 is excluded from the comparison, both main and average lateral root responses to salt correlate between our data and published data (Pearson correlation, R^2^ = 0.78 and R^2^ = 0.57 respectively, p < 0.05) (supplemental figure 3C & D). Furthermore, it is noteworthy that both main and average lateral root length responses to salt were smaller in our experiment compared to published data (supplemental figure 3). Possible explanations for these differences can be the lack of sucrose in our medium and different light sources. In conclusion, we have selected a set of accessions with reproducible variation in root system architecture responses to salt, that can be used to study the relationship between root architecture remodeling and root growth dynamics.

**Figure 1:**
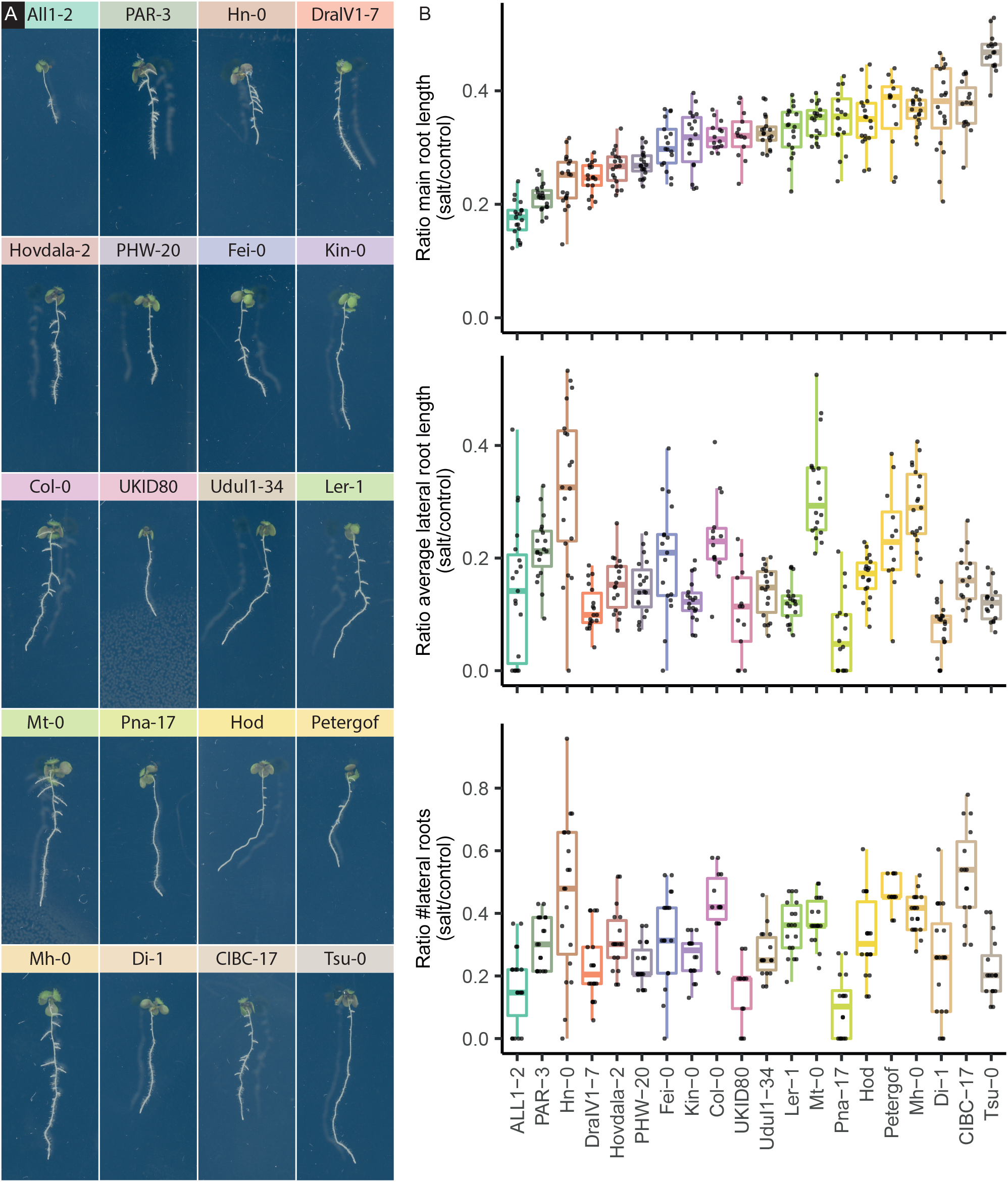
Selection of *Arabidopsis thaliana* accessions with varying root system architectures. Root architecture of 12 day-old seedlings that were transferred to 125 mM NaCl or control treatment at 5 days after sowing. A) Representative 12 day-old seedlings of the 20 different accessions under 125 mM NaCl treatment. B) Main root length, C) average root length and D) number of lateral roots quantified as ratio between NaCl treated and control. Colors correspond to headings in A. Accessions are ordered by their response in main root length to salt stress. N > 12. For this analysis outliers were removed.

It was shown before that the main root shows a multiphase growth response to high salt concentrations (Geng et al., 2013). First, we investigated if there is natural variation in these root growth dynamics. To quantify changes in main root growth phases, 7-day old seedlings were transferred to 125 mM NaCl or control plates and imaged every 20 minutes. The root growth rate varied between accessions under salt stress and control conditions (supplemental figure 4). During salt stress, we observed the previously described root growth phases, the stop, quiescent, recovery and homeostasis phase for the accession Col-0, confirming earlier studies (Geng et al., 2013). Interestingly, both the duration of these growth phases and the amount of recovery during homeostasis seem to differ between accessions (supplemental figure 4). To quantitatively compare these differences, we fitted both a single and double sigmoidal function to the root growth rate under salt stress and determined which model fitted the data better (dependent on the ΔAIC) (Figure 2, 3 and supplemental figure 5). For all accessions, except DralVI1-7, the double sigmoid fitted the growth data better, indicating that, with the exception of DralVI1-7, all accessions exhibit a decay and growth phase.

**Figure 2:**
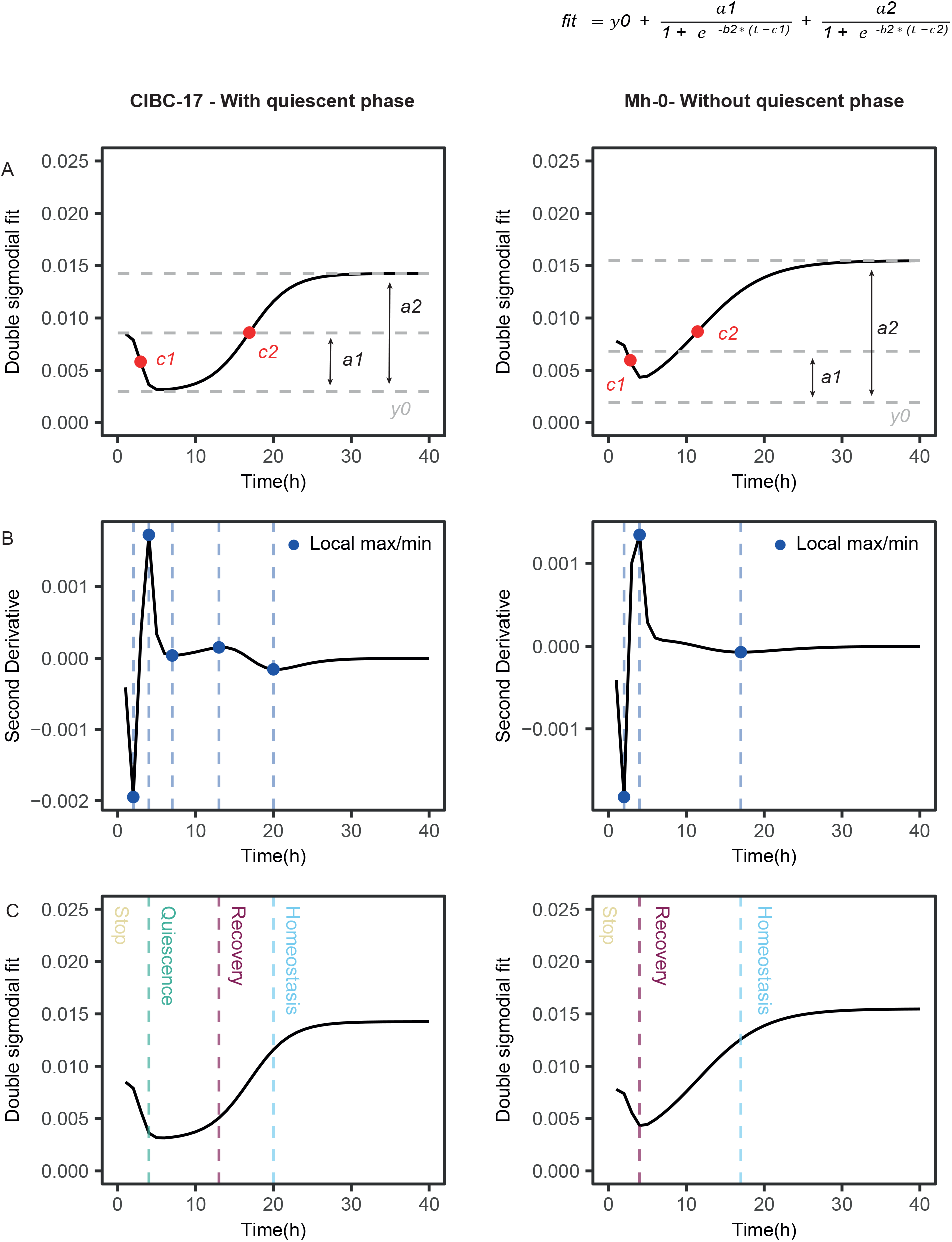
Distinguishing main root growth phases by fitting double sigmoidal equation reveals not all accessions display a quiescent growth phase. A) Fitted main root growth of 7-day old seedlings treated with 125 mM NaCl of CIBC1-7 and Mh-0 accessions. Blank line = double sigmoidal fit to main root growth data under salt stress. Red dots = first and second midpoints of the first and second sigmoid. Grey dotted lines = y0, y0 + decrease first sigmoid (a1) and y0 + increase second sigmoid (a2). B) Second derivative of the double sigmoidal fits showed in A. Blue points & blue dotted lines show the local maxima and minima of this second derivative. Accessions either show 4 (with quiescent phase) or 2 (without quiescent phase) local maxima and minima after the midpoint of the first sigmoid. C) Local maxima and minima of the second derivative were used to define the beginning and end of different growth phases. Beginning of stop, quiescent, recovery and homeostasis phase are depicted with dotted lines. A subset of accessions does not show a quiescent growth phase of the main root (without quiescent phase).

**Fig 3:**
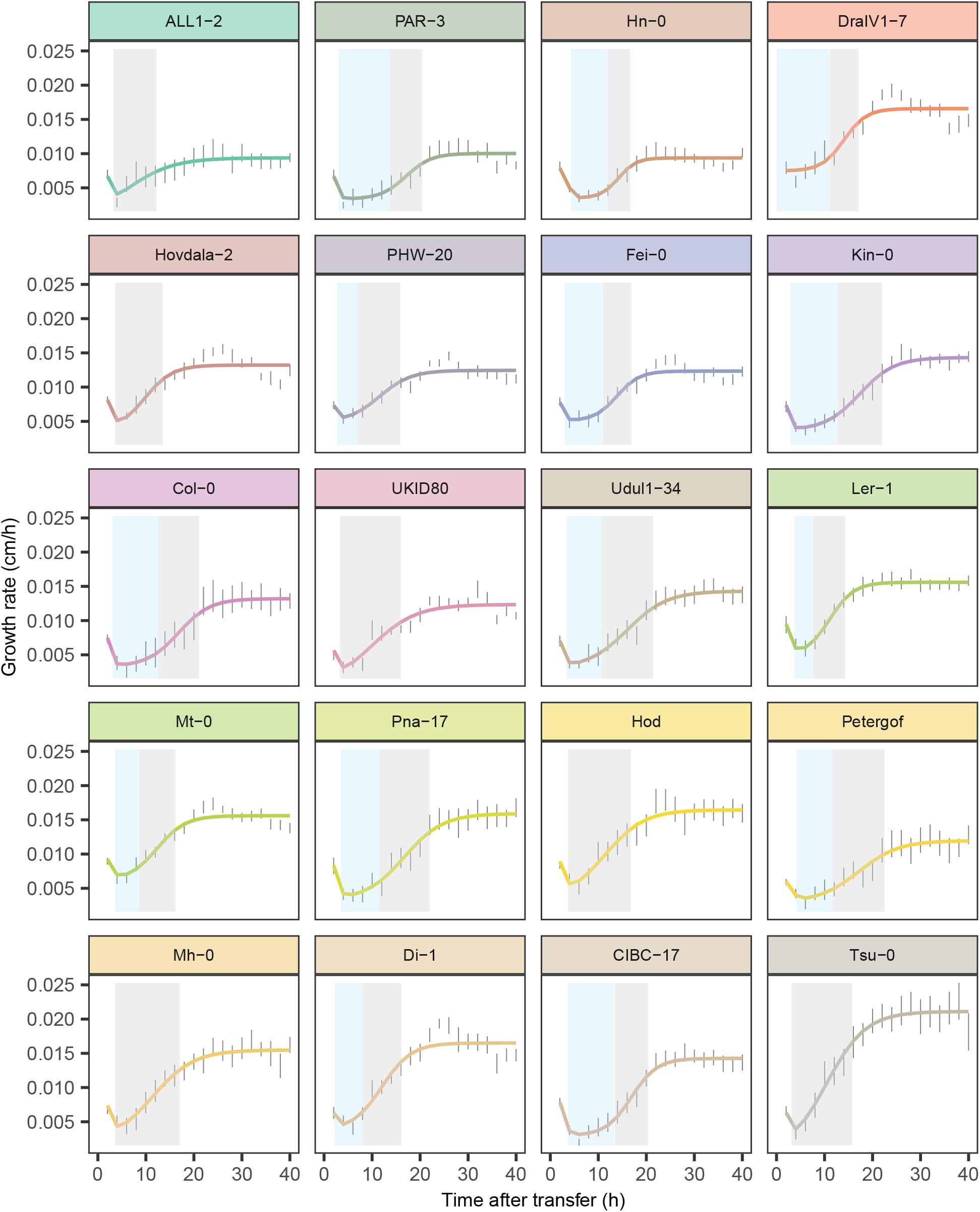
Natural variation in salt-induced main root growth phases and variation in the occurrence of a main root quiescent growth phase. Main root growth rate of 7-day old seedlings, which were transferred to 125 mM NaCl or control plates 7 days after sowing. Plates were imaged every 20 min and growth rate was averaged per hour. A double sigmoid was fitted to the growth rate over time. Accessions are depicted in subplots and ordered corresponding to their response in main root length in the overall root architecture (Figure 1B). Colored lines = double/single sigmoid fit, error bars = SE, blue shading = quiescent phase, grey shading = recovery phase, N > 8.

Subsequently, the local maxima and minima in the second derivative of the fitted models, presenting the largest changes in growth rate, were used to mark the beginning and end of separate growth phases (Figure 2). Interestingly, this analysis revealed variation in the number of root growth phases after salt stress. The main root response to salt could be grouped based on two distinct growth patterns; one with and one without a salt-induced quiescent root growth phase (Figure 2 and 3). The root growth of some accessions showed a pronounced quiescent phase (for example Di-1 and Hn-0), the growth rate of others immediately increased again after the stop phase (for example Tsu-0 and All1-2). Thus, by fitting single and double sigmoidal functions, we could quantitatively distinguish different main root growth phases under salt stress.

### Salt induces a multiphase growth response in fully developed lateral roots, and delays appearance of new lateral roots

While the main root is formed during embryogenesis, lateral roots are formed later in development from primary root tissue. This complicates measuring root growth dynamics in lateral roots, because these not only grow but are also continuously formed. A previous study measured lateral root quiescence by quantifying for how long newly emerging lateral roots showed reduced growth (Duan et al., 2013). Interestingly, salt induced a prolonged quiescent phase in these lateral roots. However, accessions of Arabidopsis also display variation in their lateral root number under control conditions (Julkowska et al., 2017). Therefore, we reasoned that the developmental stage of lateral root primordia can differ at the point of transfer between accessions, which potentially influences the timing of their quiescence. Thus, we decided to measure lateral root growth in two separate experiments; one measuring the dynamics of fully developed lateral roots and the other quantifying the effect of salt on the formation of new lateral roots.

To investigate the natural variation in lateral root growth dynamics of fully developed lateral roots, 9-day old seedlings were transferred to 125 mM salt or control plates. At this timepoint all accessions had a few apparent lateral roots. Growth rates of roots present at the moment of transfer were measured for 80 hours after transfer. While salt strongly decreased the growth rate of the main root of all accessions (supplemental figure 4), lateral root growth of some accessions (for example Pna-17) was hardly affected under salt stress (supplemental figure 6). Again, we used the single and double sigmoidal models to quantitatively describe growth phases (Figure 3 & supplemental figure 8). In this case, the double sigmoid was a better fit (determined by the ΔAIC) for only 8 accessions. Thus, in contrast to the main root, less accessions show a pronounced stop phase in the lateral root. However, all accessions show a temporal reduced growth rate in salt compared to control conditions (supplemental figure 6 & 7), which we interpreted as a quiescent phase, followed by a recovery phase. Thus, fully emerged lateral roots show similar growth phases as the main root and again not all accessions show a stop and quiescent phase. Interestingly, accessions that do not show a quiescent phase in the main root do not necessarily lack a lateral root quiescent phase and vice versa. For the accessions that show both a main and lateral root quiescent phase, the length of the main root quiescent phase is longer than that of the lateral roots, consistent with earlier reports for Col-0 (Duan et al., 2013; Geng et al., 2013).

When a seedling is turned 90° degrees (gravistimulation), the main root will follow gravity and grow down. In the bend that is formed by this turning, a local auxin maximum will synchronize the initiation of a lateral root (Lucas et al., 2008; Péret et al., 2012; Voß et al., 2015). To quantify the natural variation in lateral root development under salt stress, 5 day-old seedlings were transferred to salt and control plates and turned 90° degrees. Daily photographs of these seedlings were used to quantify the timing of lateral root appearance. Under salt stress, lateral root appearance was delayed in all accessions (Figure 5 & supplemental figure 9). More variation in lateral root appearance between accessions was observed in salt compared to control conditions. Interestingly, Pna-17 and UKID80, two accessions with the strongest delay in lateral root appearance, also show a low number of lateral roots in response to salt in the overall root architecture (Figure 1B). In conclusion, under salt stress, fully emerged lateral roots show similar growth phases as the main root and lateral root appearance after a gravistimulus is delayed compared to control conditions.

### Main and fully developed lateral roots show similar salt-induced changes in growth rate during homeostasis

Next, we investigated which of the measured growth dynamics are important for root architecture remodeling under salt and how these dynamics relate to each other. First, we calculated the salt-induced changes for the measured traits. To quantify root architecture remodeling, main root length, lateral root length and number of lateral roots were measured as a ratio between salt and control (Figure 1). Main and lateral root growth dynamics were separated in different growth phases (Figure 3 & 4), from which the length of the different phases and the average relative growth rate (salt/control) in a phase were quantified. For the accessions that did not display a certain growth phase this phase was not taken along in the analysis. Finally, we quantified the delay in lateral root appearance as a ratio between salt and control (Figure 5). This yielded a set of 16 traits measured for 20 accessions (supplemental table 2). Figure 6 shows the estimated networks among all 16 traits using the statistical model implemented in the netgwas R-package. In Figure 7A, we extracted a sub-network with respect to our traits of interest (i.e. related to the main and lateral root growth phases) from Figure 6. For the purpose of the visualizations, traits in the estimated graph are categorized with different colors based on different groups of traits that are related to: overall architecture (Figure 1), main root dynamics (Figure 3), lateral root dynamics (Figure 4) and turning assay (Figure 5). The strength of interactions is determined by calculating partial correlation values, where the thickness of edges show the magnitude of partial correlation value. Yellow and blue edges stand for positive and negative direct associations, respectively.

**Fig 4:**
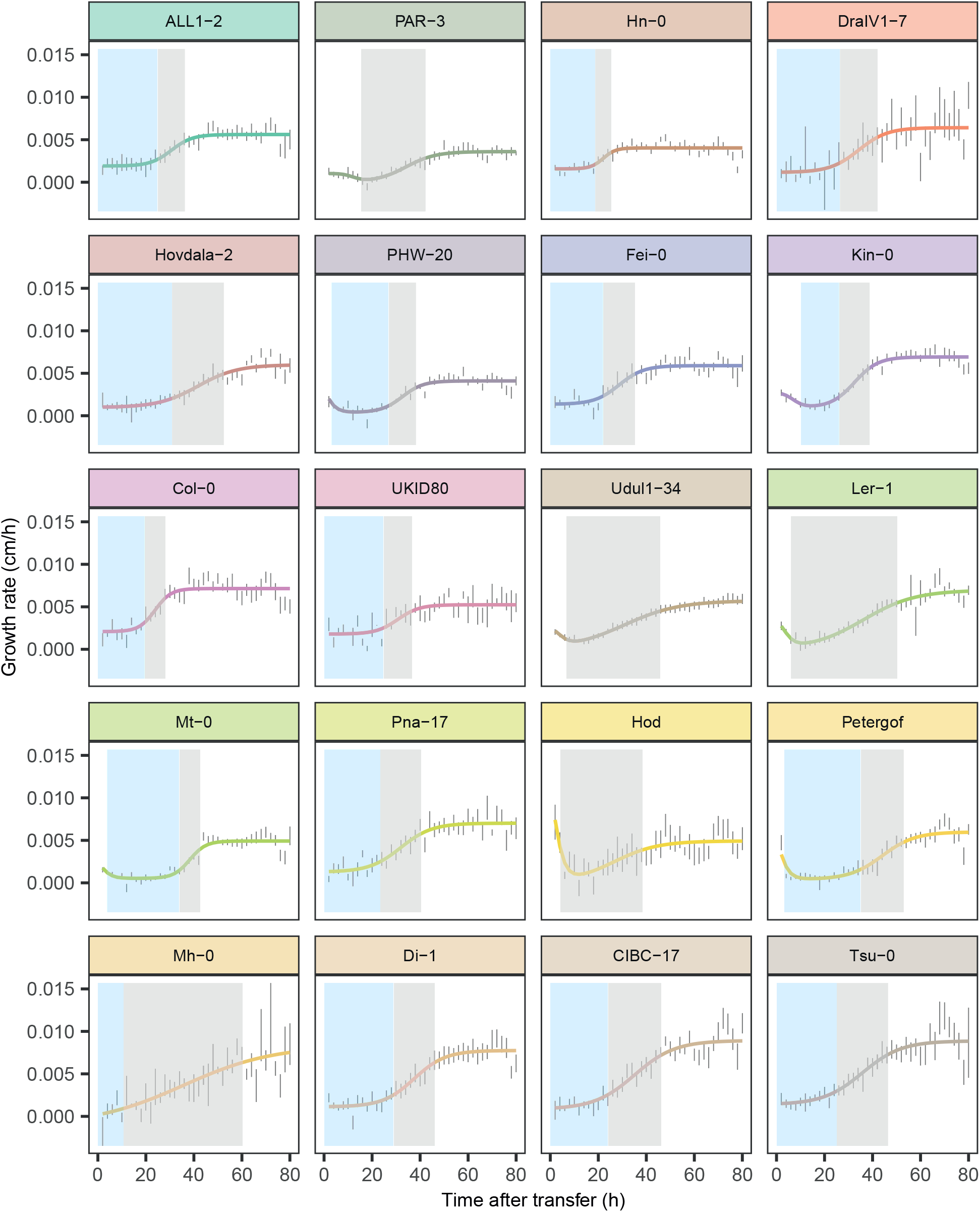
Natural variation in salt-induced lateral root growth phases and variation in the occurrence of a lateral root quiescent growth phase. Lateral root growth rate of 9-day old seedlings, which were transferred to 125 mM NaCl or control plates 9 days after sowing. The growth rate of emerged lateral roots was quantified for 80 hours after transfer. A double sigmoid was fitted to the growth rate over time. Accessions are depicted in subplots and ordered corresponding to their response in main root length in the overall root architecture (Figure 1B). Colored lines = double/single sigmoid fit, error bars = SE, blue shading = quiescent phase, grey shading = recovery phase, N > 3.

**Figure 5:**
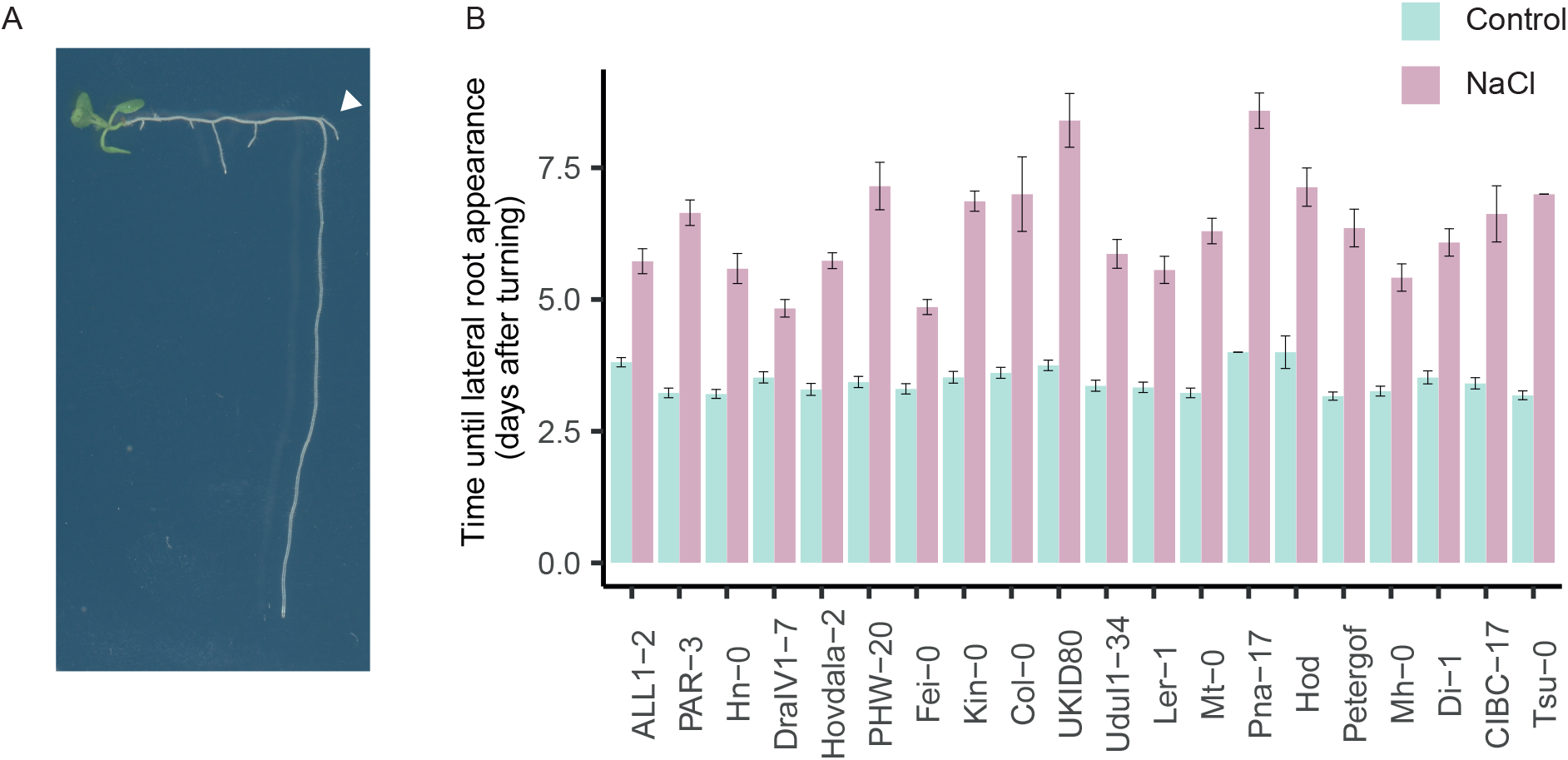
Salt induces a delay in lateral root appearance in gravistimulated seedlings. Seedlings were transferred to 125 mM NaCl or control plates and turned 90° degrees. Plates were photographed daily and timing of appearance of a lateral root in the bending zone was determined. A) Example image of a gravistimulated seedling under control conditions. White arrowhead = lateral root in the bending zone. B) Time until lateral root appearance under NaCl and control treatment in different accessions. Bars = mean of accession, error bar = SE, for N per genotype see supplemental figure 9.

**Figure 6:**
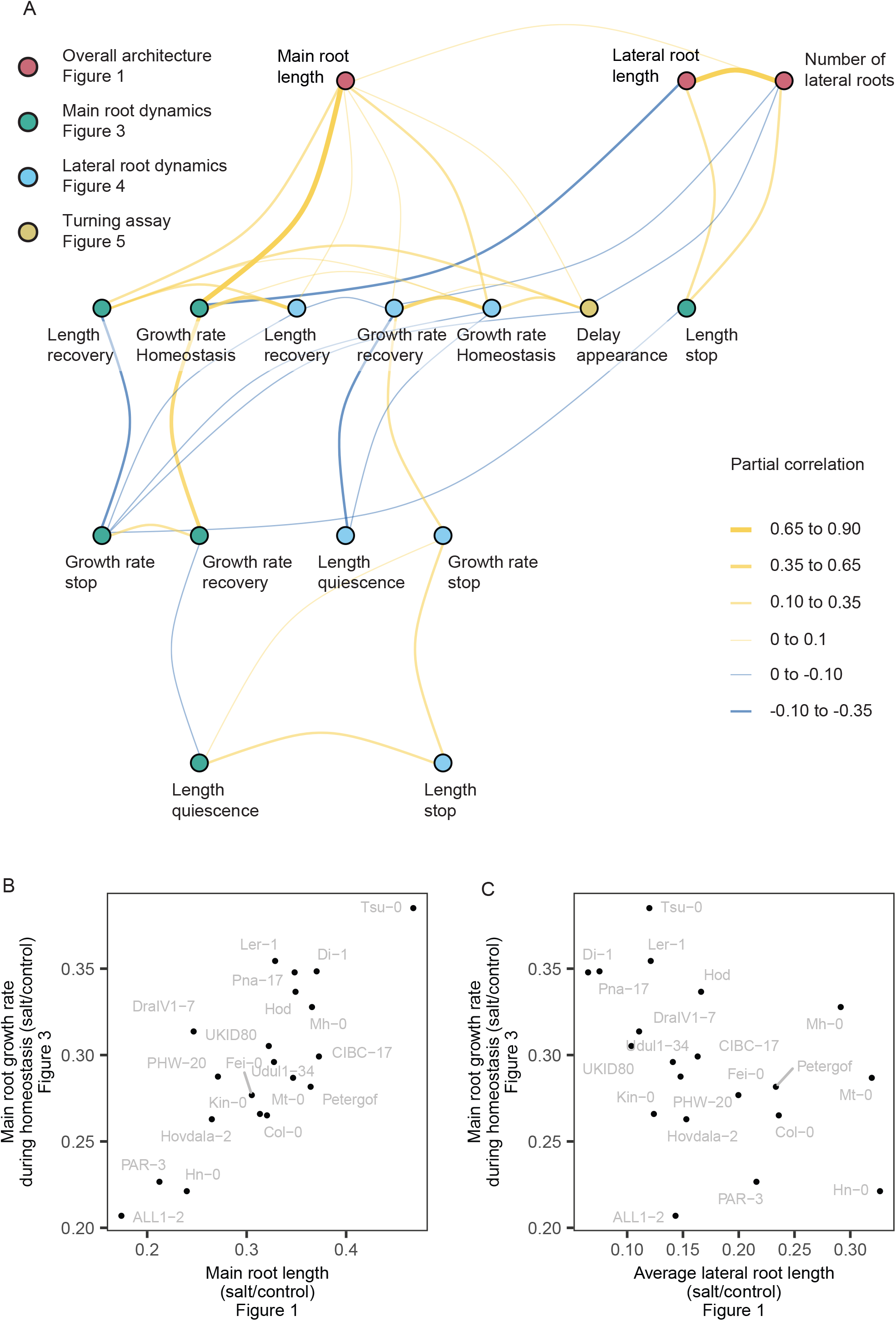
The speed of the main root during homeostasis shows a trade-off between responses in main and lateral root length under salt stress. A) Partial correlation network between overall root architecture traits (Figure 1) and dynamic root growth traits (Figure 3, 4 and 5). Nodes depict traits and are colored depending on the experiment in which they were measured. Edges depict partial correlations and are colored depending on the strength of the correlation (yellow = positive, blue = negative). All traits, except for the length of the phases, are depicted as a ratio between salt and control conditions. B) Scatter between response in main root length to salt in the overall root architecture (Figure 1) and relative growth rate of the main root during homeostasis in salt stress (Figure 3). C) Scatter between response in average lateral root length to salt in the overall root architecture (Figure 1) and relative growth rate of the main root during homeostasis in salt stress (Figure 3).

We showed that main roots and fully developed lateral roots display similar root growth dynamics under salt stress (Figure 2 and 3). First, we investigated if the growth dynamics during salt stress are correlated between main and lateral roots amongst accessions using a subset of the estimated network (Figure 7A). Between main and lateral roots, the length of the recovery phase and the growth rate in during homeostasis are positively conditionally correlated among accessions (Figure 7A). For example, accessions with a longer main root recovery phase also show a longer lateral root recovery phase (Figure 7B). This indicates that main and lateral roots of these accessions show similar behavior under salt stress in later growth phases. Intriguingly, we observed no direct positive correlation between the length of the main and lateral root quiescent phases (Figure 7A & C). Moreover, accessions lacking a main root quiescent phase generally do show a lateral root quiescent phase and the other way around, except for Hod (Figure 7C). Thus, accessions show different relative lengths of the quiescent phase between main and lateral roots. Both the length of the quiescent phase and the length of the recovery phase are in general longer for the lateral roots compared to the main root (Figure 7B & C). In conclusion, even though main and lateral roots respond similarly to salt in the recovery and homeostasis phase, the length of the salt-induced quiescent phase is not consistent between these tissues.

**Figure 7:**
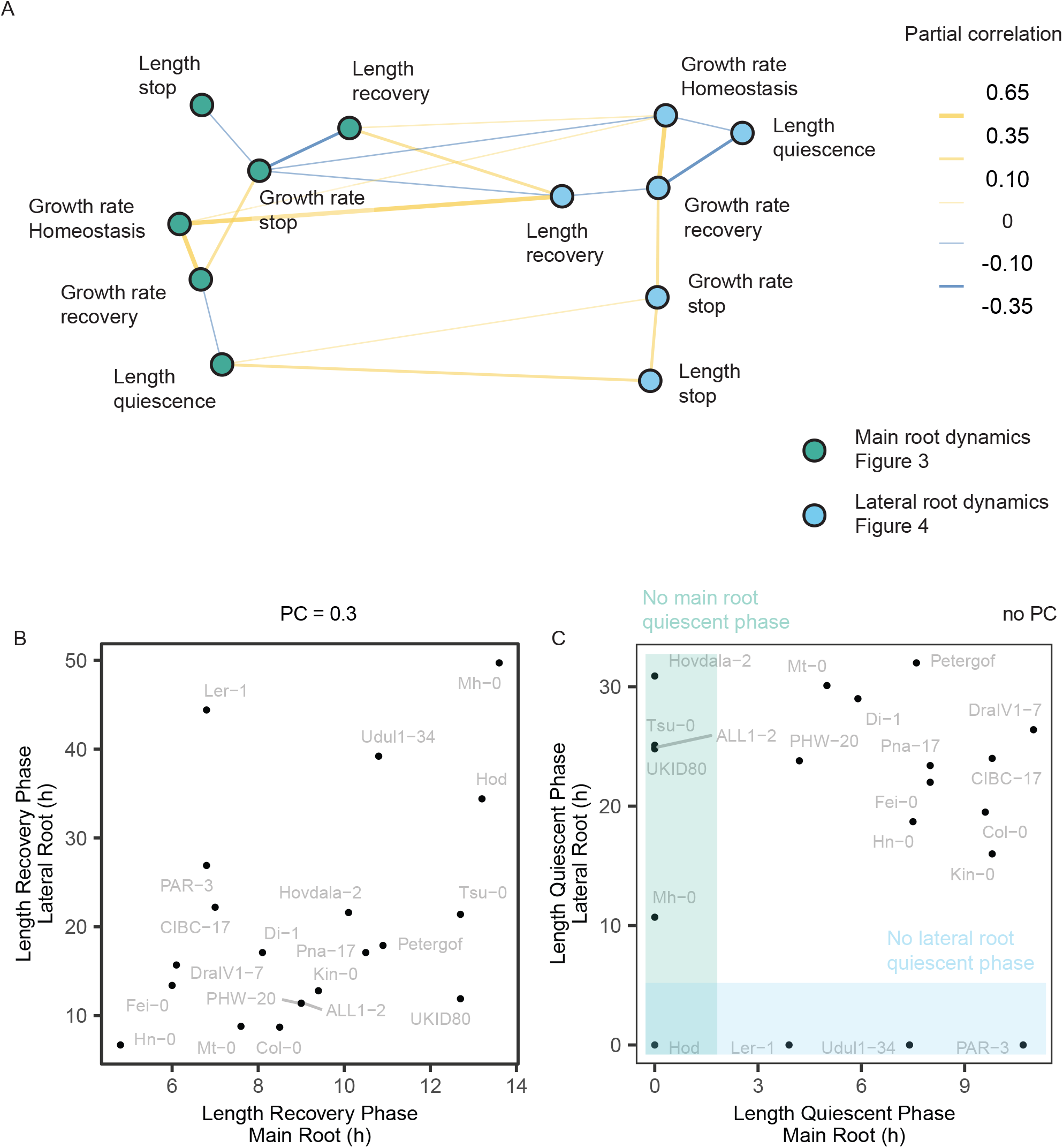
The length of the recovery phase and growth rate during homeostasis positively correlate between main and lateral roots. A) Partial correlation network between main (Figure 3) and lateral (Figure 4) root growth phases (this is a subset of the network shown in figure 7). Green and blue nodes = length and speed in growth phases in main and lateral roots, respectively. Speed = average relative growth rate (salt/control) within the timeframe of a growth phase. Edges = partial correlation between traits (yellow = positive, blue = negative, width and shade show the strength of the correlation). B) Scatterplot between length of the main and lateral root recovery phase. C) Scatter plot between the length of the main and lateral root quiescent phase. Green shading = accessions without a main root quiescent phase. Blue shading = accessions without lateral root quiescent phase.

### The main root growth rate during homeostasis and lateral root appearance reveal trade-offs between investing in main or lateral root tissue under salt stress

Next, we investigated which dynamic root traits are correlated to the changes in overall root architecture during salt (Figure 6). Among traits of the overall root architecture, the salt-induced changes in lateral root number and lateral root length showed a strong direct positive correlation. There was a weaker direct correlation between the relative in main root length and relative lateral root number under salt stress. In contrast, the changes in main root length were not directly correlated to the changes in average lateral root length under salt stress. The relative main root length in the overall root architecture showed a positive partial correlation with the relative main root growth rate during the homeostasis and the length of recovery phase (Figure 6B). Thus, a fast growing main root leads to a longer main root in the overall architecture and a longer recovery phase might be necessary to maintain main root length. Interestingly, the relative main root growth rate during homeostasis also directly but negatively correlated with the relative average lateral root length, suggesting the existence of a trade-off between a fast growing main root and the investment in lateral root elongation under salt (Figure 6C). This trade-off cannot be explained by the lateral root growth rate during homeostasis, because this trait is directly positively correlated to the main root growth rate during homeostasis and to the main root length. Thus, lateral root growth rate during homeostasis appears less important for root architecture remodeling than main root growth rate during homeostasis.

The changes in lateral root number negatively correlate to the relative lateral root appearance in the turning assay. Thus, accessions that show a long salt-induced delay in lateral root appearance also end up with less lateral roots in their root architecture. Furthermore, the delay in lateral root appearance also directly positively correlates to the main root length, suggesting that accessions with a relatively longer main root during salt stress generally also show a long salt-induced delay in their lateral root appearance in a turning assay. Thus lateral root appearance is another trait that could involve in inducing a trade-off between investing in main or lateral root tissue. This makes lateral root appearance an interesting trait to potentially predict root architecture remodeling. Both the salt-induced changes in average lateral root length and lateral root number relate to the length of the stop phase of the main root. This connection could indicate that an early salt-induced mechanism influences both the reduction in main root growth rate and initiation and outgrowth of lateral roots. Neither length of the main or lateral root quiescent phases are directly correlated to the overall root architecture, but they both negatively correlate to the growth rate in the recovery. Thus, we do not find a direct link between the length of root quiescence and root architecture remodeling, but both the main root growth rate during homeostasis and lateral root appearance reveal possible trade-offs for investing in either main or lateral root mass during salt stress.

## Discussion

In Col-0, salt induces a multiphase root growth response in the main root and prolonged lateral root quiescence of the lateral roots (Duan et al., 2013; Geng et al., 2013). The quiescent phase of the lateral roots occurs at lower salt concentrations and is longer than that of the main root (Duan et al., 2013; Geng et al., 2013). For both main and lateral roots, we found natural variation in the length of and growth rate during salt-induced growth phases and in the occurrence of a salt-induced quiescent phase. We can confirm that the quiescent phase, but also the recovery phase, of lateral roots is generally longer than in the main root and this holds true for all tested accessions that show both a main and lateral root quiescent phase. Between main and lateral roots, there is a positive correlation for the length of the recovery phase and the growth rate during homeostasis. Thus, in these later growth phases main and lateral roots show a similar response to salt stress and these phases could potentially be regulated through a common pathway in both tissues. In contrast, the length of the quiescent phase is not correlated between main and lateral roots and, apart from Hod, accessions express either a lateral or main root quiescent phase, but not both. Interestingly, abscisic acid (ABA) is known to regulate both main and lateral root growth dynamics, but in a contrasting fashion (Duan et al., 2013; Geng et al., 2013). In the main root, ABA promotes growth, and increased ABA signaling results in a short main root quiescent phase (Geng et al., 2013). In contrast, endodermal ABA signaling is necessary to induce lateral root quiescence in the lateral roots (Duan et al., 2013). Furthermore, accessions have been shown to exhibit natural variation in root growth sensitivity in response to ABA and these differences could be linked to their strategy in modulating root architecture in response to salt (Julkowska et al., 2014). The dual role of ABA in promoting main root growth but inhibiting lateral root growth, makes it an interesting hormone to potentially induce differential responses in salt-induced growth quiescence of the main and lateral roots.

Salt stress alters both cell elongation and cell division (West et al., 2004). In the main root, the activity of cyclin-dependent kinases (CDKs), which are necessary for cell cycle progression, is temporarily reduced under salt stress (West et al., 2004), coinciding with the salt-induced growth quiescence of the main root. In emerged lateral roots, the activity of the cell cycle reporter *CYCB1:1:GUS* was downregulated under salt stress compared to control conditions (Duan et al., 2013). It is possible that the quiescent phase is caused by a reduction of cell cycle activity in the meristem. Interestingly, a reduction in cell cycle genes and cell division under salt stress was shown to be conserved in the halophyte (salt-tolerant) species *Schrenkiella parvula*. However, under 125 mM of salt stress, an increase in cell elongation compensates for this reduction in cell division and salt increases the main root growth rate in this species (Li et al., 2022). We find natural variation in the length of the main and lateral root growth quiescence phase under salt stress, however these differences do not directly correlate to differences in overall root architecture. Further research is needed to investigate the potential role of the quiescent phase in adaptation to salt stress.

Our data showed that the relative main root length under salt stress positively correlates with the length of the recovery phase and the growth rate during homeostasis. In contrast, lateral root length does not correlate to the length of or speed during the recovery or homeostasis phase. This can possibly be due to the age at which the plants were transferred (5 day-old) and the age at which overall root-architecture was quantified (12 day-old). At 5 day-old, seedlings do not have emerged lateral roots yet, but we measure the growth phases in emerged lateral roots. Therefore it could be that the growth phases of the lateral roots contribute to root architecture when the plants are transferred later. At 12 day-old, salt treated plants still have relatively small lateral roots (Figure 1A), which could make the time at which lateral roots are formed more important for their final length than the growth rate. In agreement with this, the salt-response in lateral root length in the overall root architecture shows a strong positive correlation with the salt-response in lateral root number. Alternatively, the low lateral root growth rate together with the long quiescent phase and delay in lateral root appearance make the lateral root growth rate during homeostasis less important for overall lateral root length.

In control conditions, main and lateral root length are positively correlated, but during salt stress this correlation is lost, suggesting a salt-induced trade-off between main and lateral root length (Julkowska et al., 2014). A possible explanation for this trade-off could be that under salt stress the energy available for root growth is limiting, due to inhibition of photosynthesis and reduced water uptake. Under these conditions, accessions could either invest in main or lateral root growth, inducing a trade-off. This hypothesis aligns with the observation that accessions employ different strategies in modifying root growth, partly dependent on relative investment in main or lateral root tissue (Julkowska et al., 2014). It would be interesting to investigate if different root architecture strategies under salt stress could be beneficial depending on, for example, the soil type and distribution of salt in the soil. Our data suggest that the relative growth rate of the main root during homeostasis and the delay in lateral root appearance under salt influence this trade-off, because they are oppositely correlated (positive vs negative) between main and lateral root traits in the overall root architecture. We show that main root length is influenced by the length of the recovery phase and the growth rate during homeostasis, while lateral root traits are partly influenced by the delay in lateral root appearance. These traits might be interesting to predict root system architecture. After investigating what type of root system is optimal in response salt, the discovery of genes and biological processes regulating these traits could provide a toolset to modulate root architecture to improve performance under salt stress.

In conclusion, we show that main and lateral root growth responses to salt are linked during growth recovery and homeostasis. However, the length of growth quiescence does not correlate between main and lateral roots. By correlating growth dynamics to overall root architecture, we show that the length of the main root recovery phase and the main root growth rate during homeostasis are important for the remodeling of overall main root length. In contrast, the salt-induced delay in lateral root appearance partly explains the salt-induced changes in lateral root traits. Both the main root growth rate during homeostasis and the delay in lateral root appearance reveal an interesting trade-off between investing in main and lateral root length in response to salt.

## Acknowledgements

Thanks to Yutao Zou for providing the *Arabidopsis* accessions. Thanks to Jasper Engel for providing some guidance with model fittings. This work is supported by the European Research Council (ERC) under the EU Horizon 2020 Research and Innovation program (grant agreement 724321; Sense2SurviveSalt) to C.T.

## Author contributions

EvZ, CT, CG and PB planned and designed the research. EvZ, EJM and SBA carried out experiments. EvZ, SBA and PB analyzed the data. EvZ, CT and CG wrote the manuscript. All authors read and provided feedback on the manuscript.

## Figure captions

**Supplemental figure 1:**
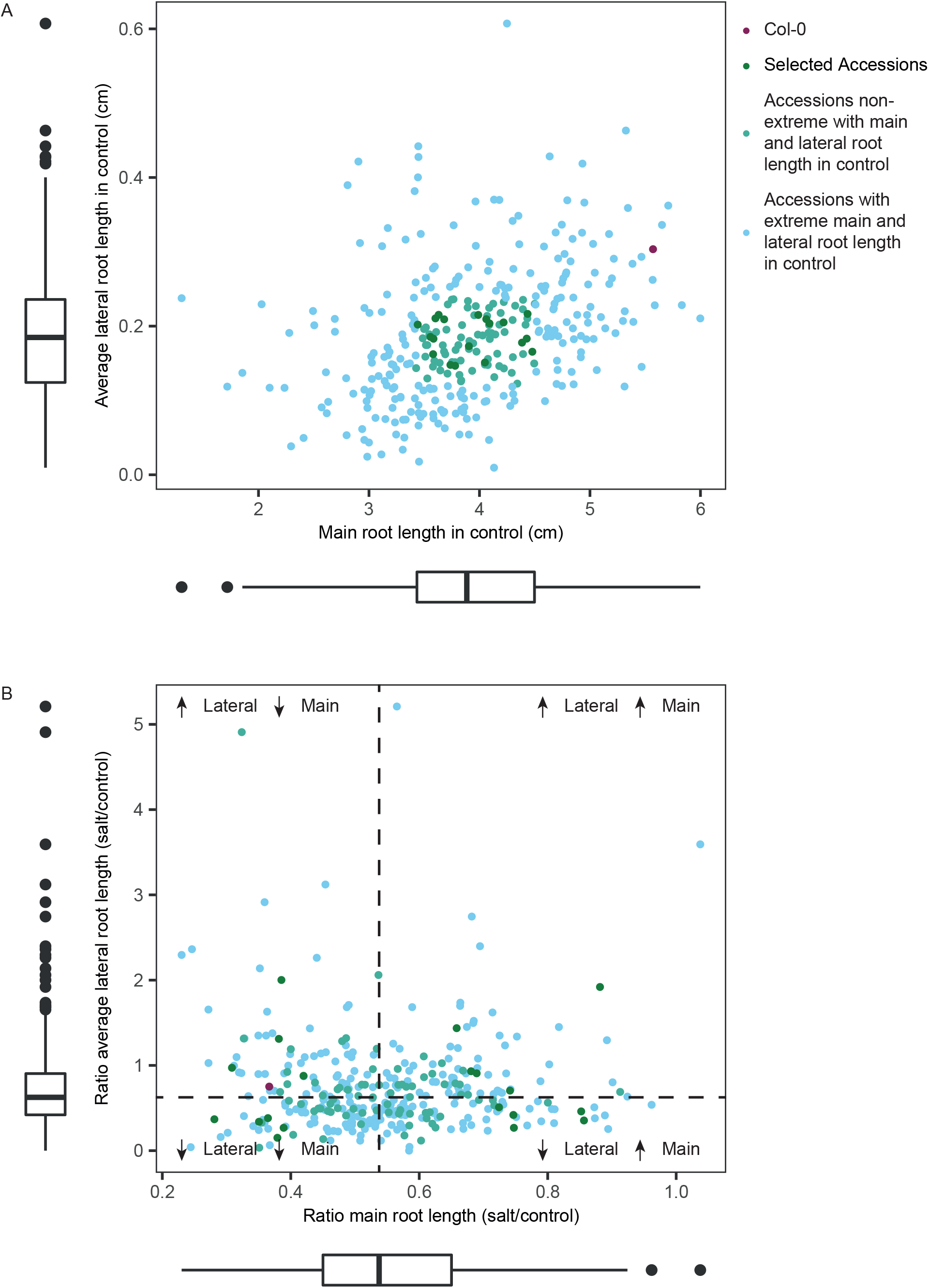
Selection of a subset of accessions with variable root system architecture responses to salt. Publicly available data (Julkowska et al., 2017) was used to select a set of 20 accessions with variable root architecture responses to salt but without extreme main or average lateral root length in control conditions. A) Scatter of main and average lateral root length of different accessions in control conditions. Accessions within the interquartile range (+25% and -25% from median, box plots plotted alongside axes) of both main and average lateral root length were determined to have normal architectures under control conditions. B) Scatter of the changes in main and average lateral root length during salt stress for different accessions, expressed as a ratio between salt and control. From the accessions without an extreme main root and lateral root length in control conditions, a set of 20 accessions were selected that showed varying responses in main and/or average lateral root length (dark green). Col-0 was added although it did show high main and average lateral root length in control conditions. Blue = accessions with extreme root architecture in control, light green = accessions with average root architecture in control, dark green = selected accessions, red = Col-0. Dotted lines represent the median as shown in the box plots, and serve to divide the accessions based on high or low average main or lateral root growth.

**Supplemental figure 2:**
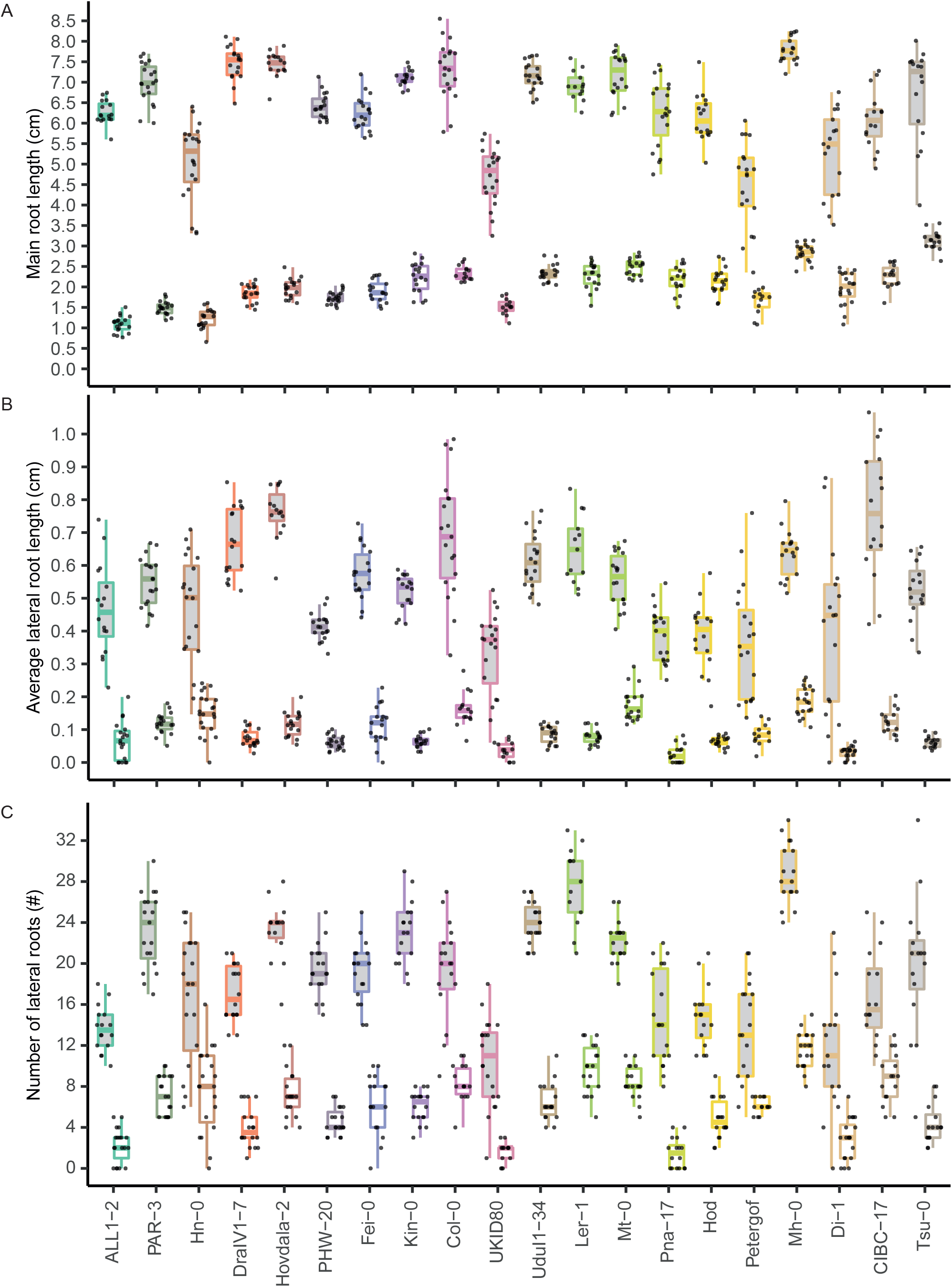
Root architecture traits of selection of *Arabisopsis thaliana* accessions. Root architecture of 12 day-old seedlings that were transferred to 125 mM NaCl or control treatment at 5 days after sowing. A) Main root length, B) average lateral root length and C) number of lateral roots of a selection of 20 accessions under 125 mM NaCl (white boxes) and control (grey boxes) treatment. Accessions are ordered per main rot length responses and colors per accession corresponding to Figure 1. N > 12. For this analysis outliers were removed.

**Supplemental figure 3:**
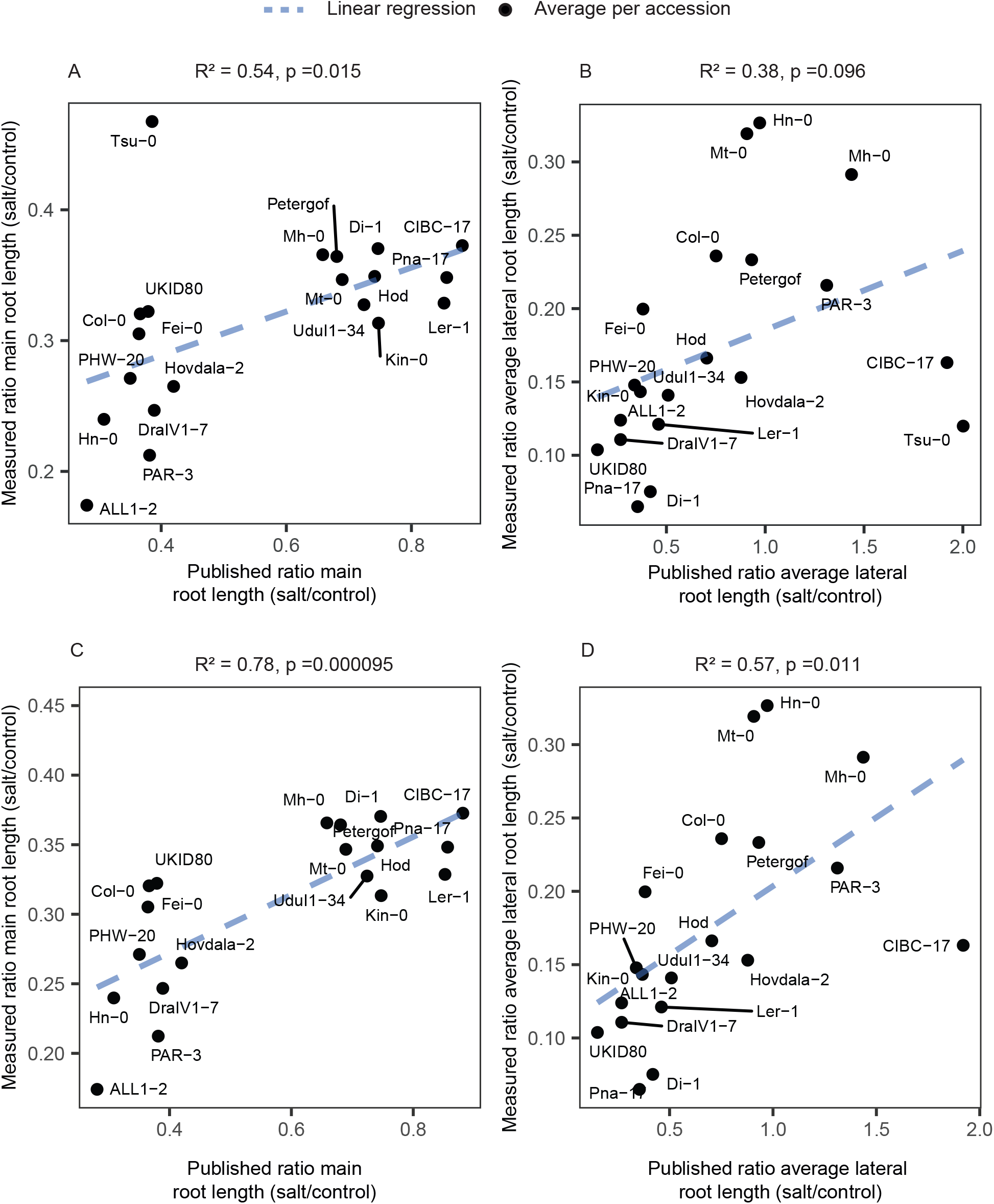
Main root length changes during salt stress are correlated between experiments, but average lateral root length responses to salt are not. Scatter plots of A) main root length responses to salt and B) average lateral root length responses to salt comparing averages per accession as measured in this study (y axis) with those measured in previous study (x axis) (Julkowska et al., 2017). Dots = average per accession, blue dotted line = linear regression, R = R squared, p = p-value of the regression.

**Supplemental figure 4:**
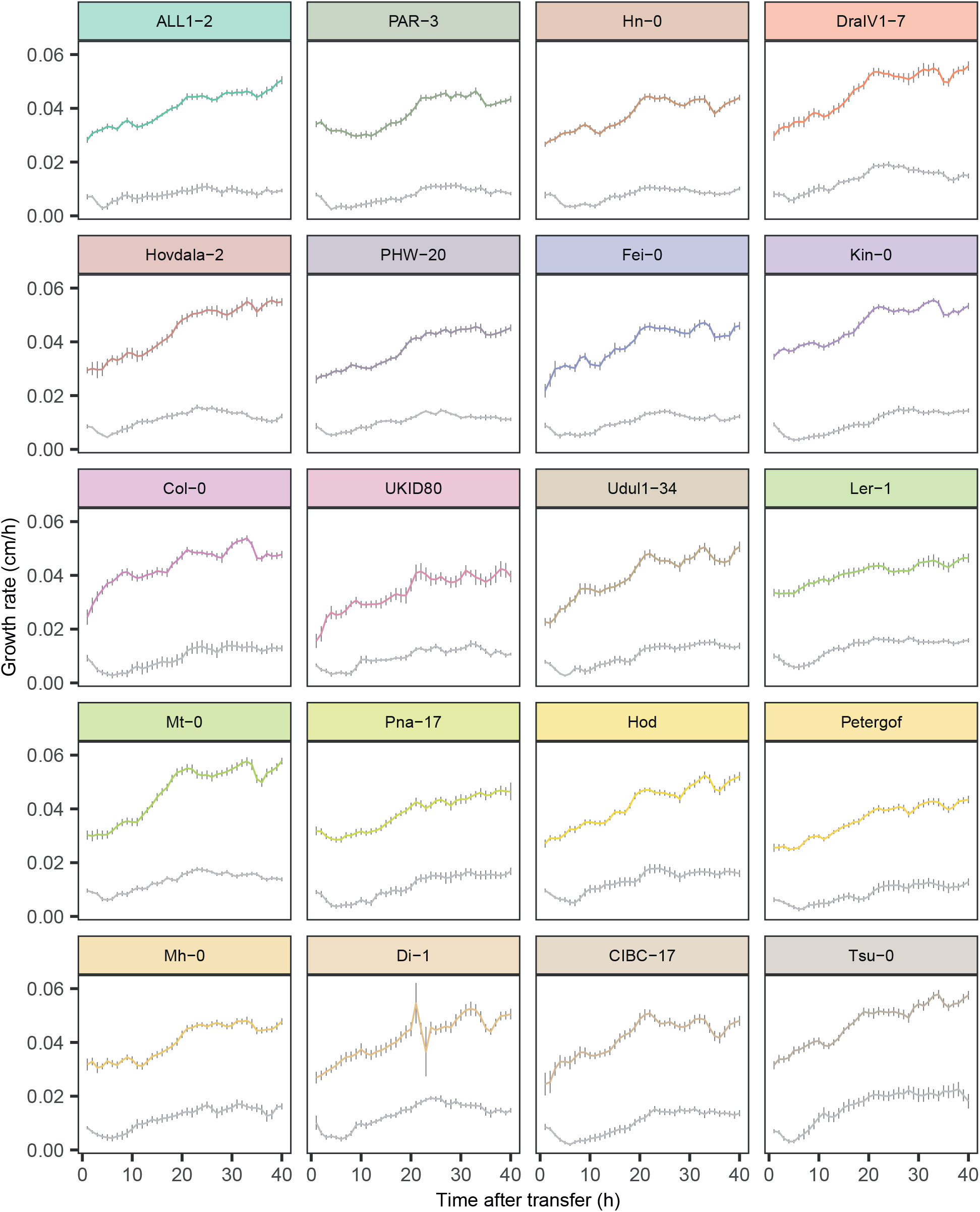
Growth rate of the main root under salt and control conditions in 20 accessions of *Arabidopsis thaliana*. Growth rate of the main root of seedlings that were transferred to 125 mM NaCl or control plates at 7 day-old. Plates were imaged every 20 min and growth rate was averaged per hour. Accessions are depicted in subplots and ordered corresponding to their response in main root length in the overall root architecture (Figure 1B). Colored line = control treatment, grey line = salt treatment, error bars = SE, N > 8.

**Supplemental figure 5:**
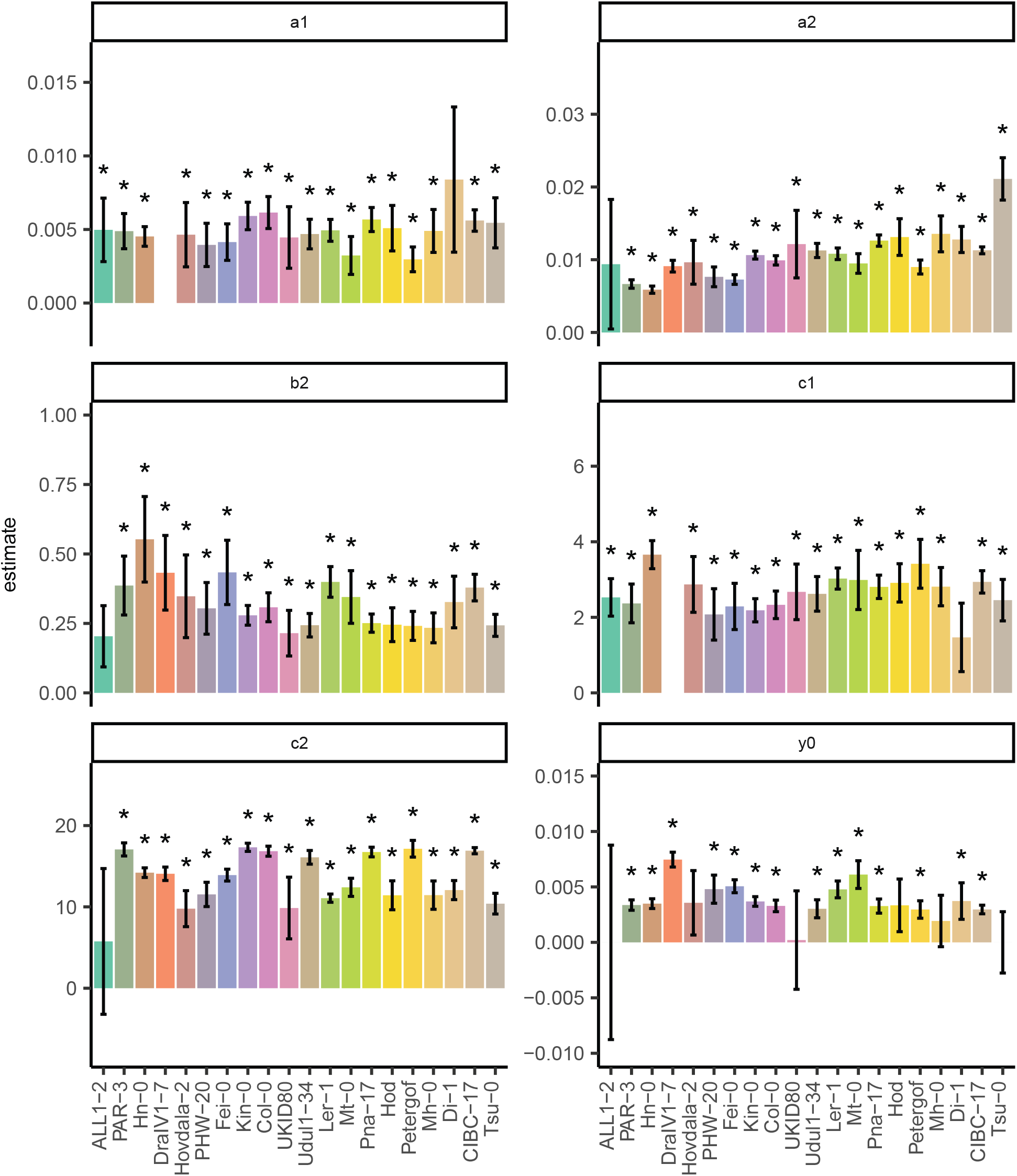
Parameter estimates for double sigmoidal that was fitted to main root growth rate under salt stress. A double sigmoidal equation was fitted to the main root growth rate under salt stress using nlsLM package. Estimates of the different parameters in the model are shown in subplots for the main root growth of 20 accessions. Accessions are ordered corresponding to their response in main root length in the overall root architecture (Figure 1B). Errorbars = SE of the estimation, * = significant parameter estimate (nlsLM, p < 0.05).

**Supplemental figure 6:**
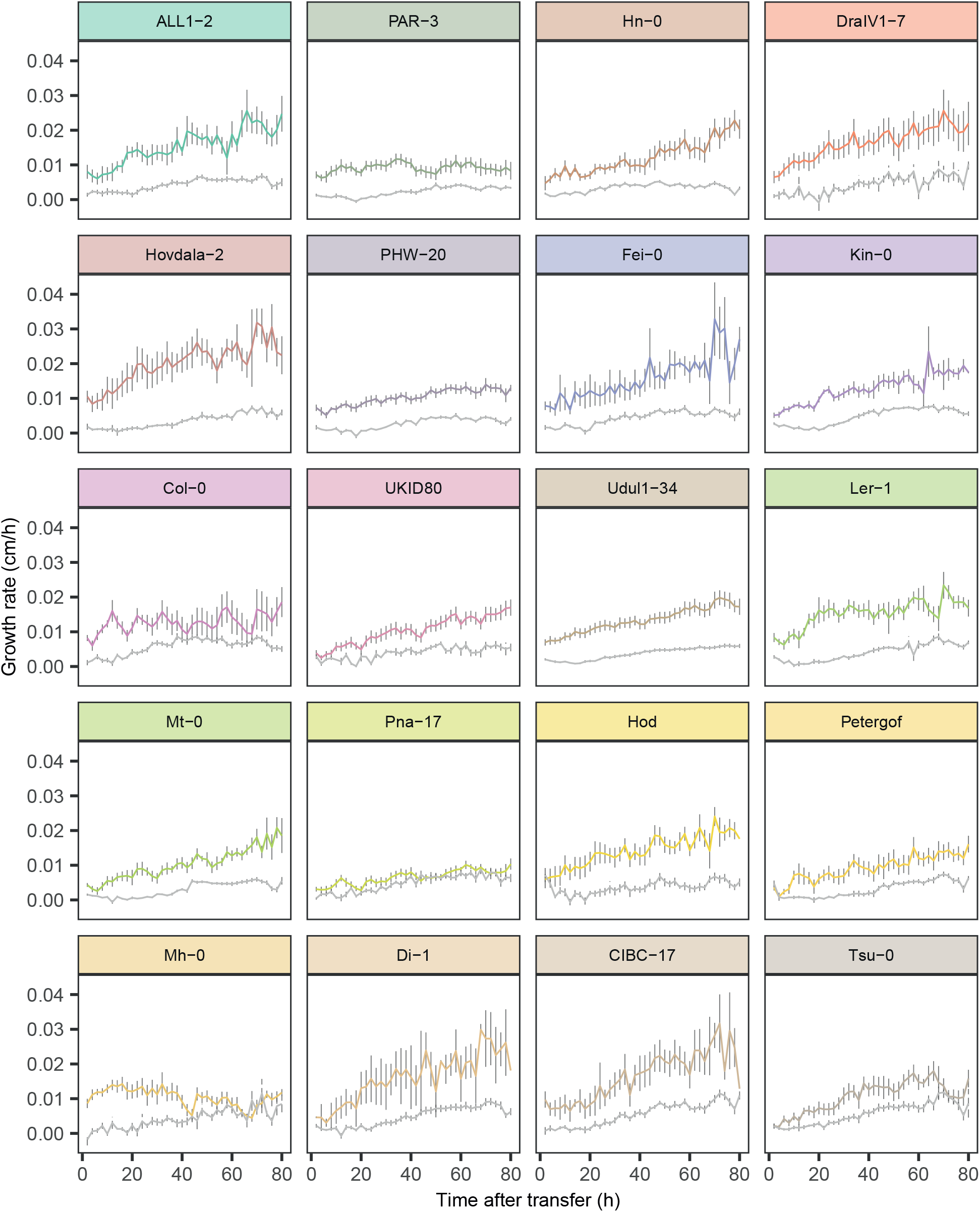
Growth rate of the emerged lateral roots under salt and control conditions in 20 accessions of *Arabidopsis thaliana*. Growth rate of the lateral roots of seedlings that were transferred to 125 mM NaCl or control plates at 7 day-old. Only lateral roots that were present on the moment of transfer were included in the analysis. Plates were imaged every 20 min and growth rate was averaged per hour. Accessions are depicted in subplots and ordered corresponding to their response in main root length in the overall root architecture (Figure 1B). Colored line = control treatment, grey line = salt treatment, error bars = SE, N > 2.

**Supplemental figure 7:**
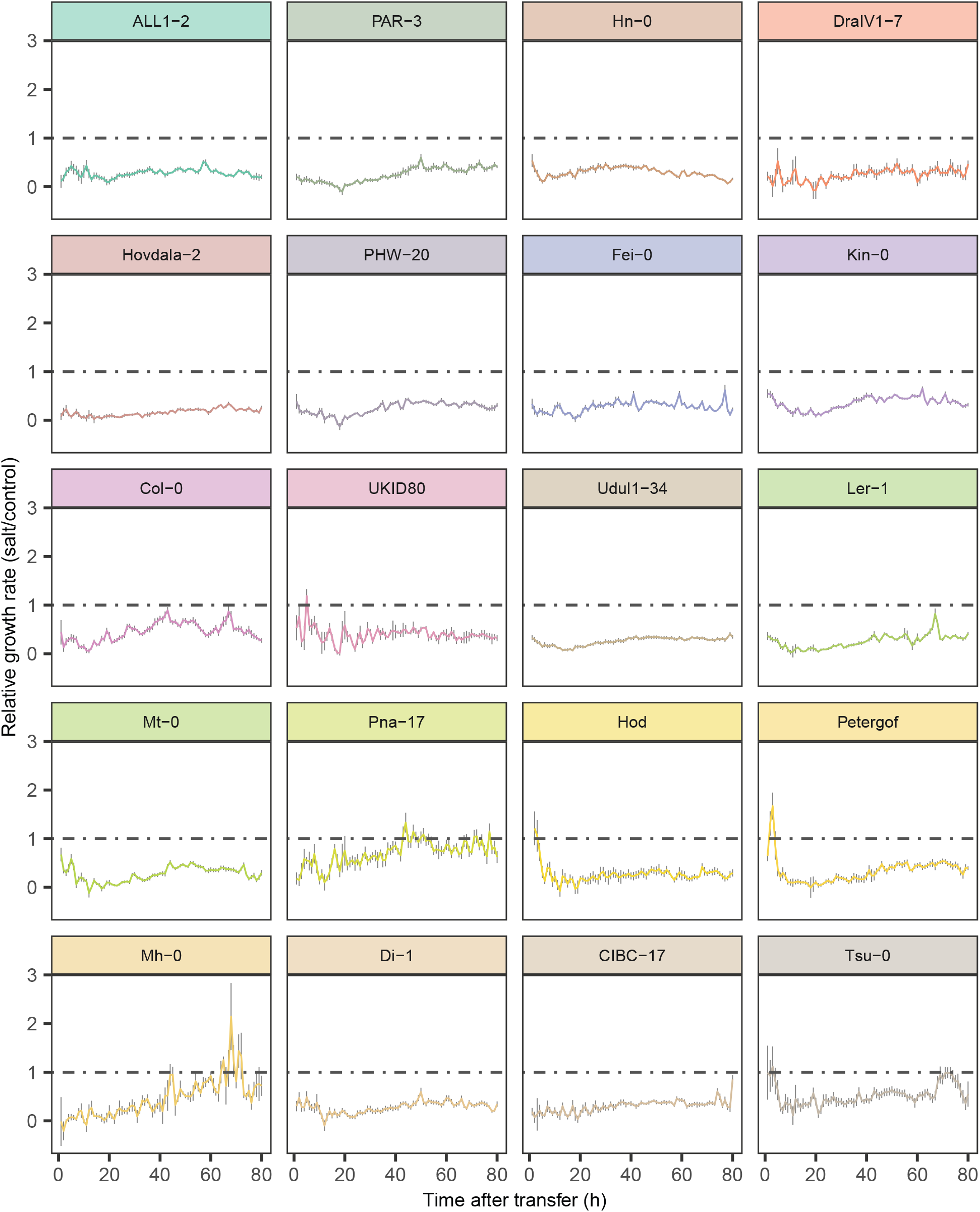
Relative lateral root growth of emerged lateral roots under salt and control conditions in 20 accessions of *Arabidopsis thaliana*. Seedlings were transferred to 125 mM NaCl or control plates at 7 day-old and the lateral root length was measured every 20 min. Only lateral roots that were present on the moment of transfer were included in the analysis. Growth rate was averaged per hour and divided by the growth rate in control conditions at the same timepoint. Accessions are depicted in subplots and ordered corresponding to their response in main root length in the overall root architecture (Figure 1B). Colored line = relative growth rate in salt treatment compared to control conditions, dotted grey line = ratio of 1, error bars = SE, N > 2.

**Supplemental figure 8:**
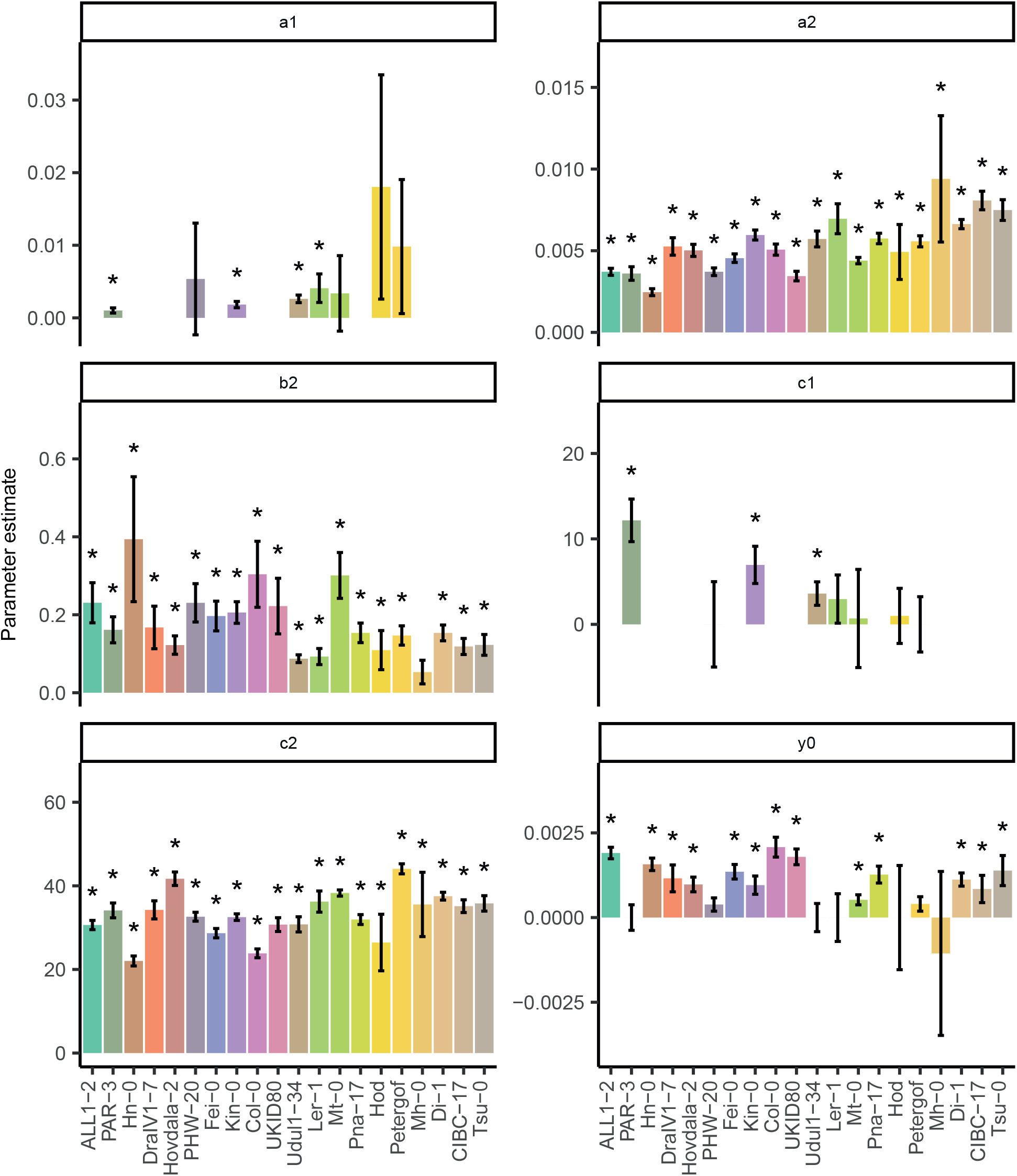
Parameter estimates for double or single sigmoid that were fitted to lateral root growth rate under salt stress. Both a double and single sigmoidal equation were fitted to the lateral root growth rate under salt stress using nlsLM package. The double sigmoid was used as a fit if the AIC of the double sigmoid was 2 or more higher than the AIC of the single sigmoid. For the lateral root growth of 20 accessions, estimates of the different parameters of the best fitting model are shown in subplots. For the parameters where a single sigmoid was the best fit a1 and c1 are lacking. Accessions are ordered corresponding to their response in main root length in the overall root architecture (Figure 1B). Errorbars = SE of the estimation, * = significant parameter estimate (nlsLM, p < 0.05).

**Supplemental figure 9:**
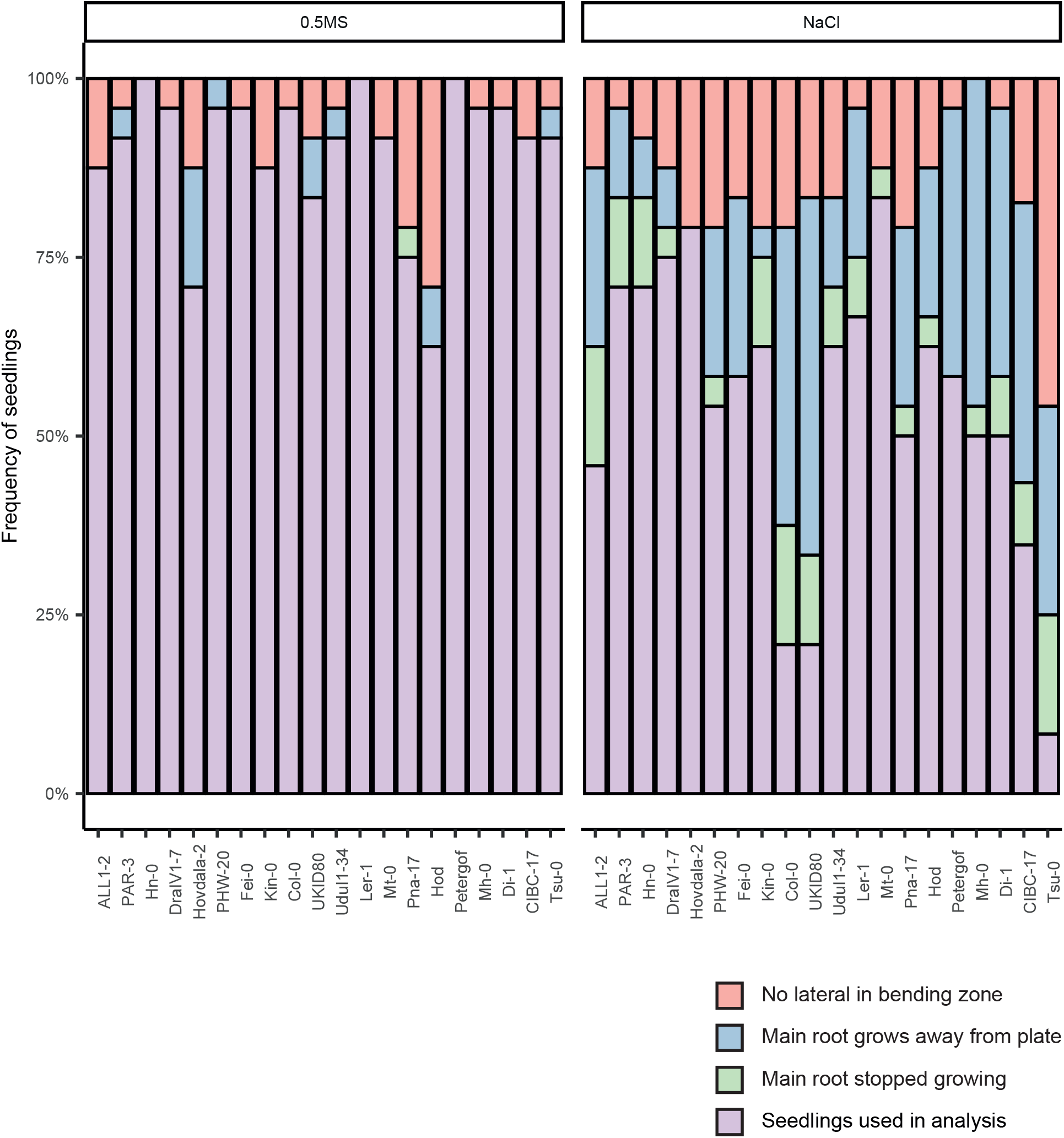
Proportion of seedlings that were taken out of the analysis of the turning assay. Seedlings were transferred to 125 mM NaCl or control plates and turned 90° degrees. The timing of appearance of a lateral root in the bending zone was determined. In purple the proportion of seedlings that was included in the analysis is depicted. Seedlings were excluded for not showing a lateral root in the bending zone (red), the main root growing away from the plate (blue) or the main root not continuing growing after transfer (green), N = 24.

## Literature

Awlia M, Alshareef N, Saber N, Korte A, Oakey H, Panzarová K, Trtílek M, Negrão S, Tester M, Julkowska MM (2021) Genetic mapping of the early responses to salt stress in Arabidopsis thaliana. Plant J 107: 544–563

Awlia M, Nigro A, Fajkus J, Schmoeckel SM, Negrão S, Santelia D, Trtílek M, Tester M, Julkowska MM, Panzarová K (2016) High-throughput non-destructive phenotyping of traits that contribute to salinity tolerance in Arabidopsis thaliana. Front Plant Sci 7: 1–15

Baione F, Biancalana D, De Angelis P (2020) An application of Sigmoid and Double-Sigmoid functions for dynamic policyholder behaviour. Decis Econ Financ 44: 5–22

Behrouzi P, Wit EC (2019) Detecting epistatic selection with partially observed genotype data by using copula graphical models. J R Stat Soc Ser C Appl Stat 68: 141–160

Caglar MU, Teufel AI, Wilke CO (2018) Sicegar: R package for sigmoidal and double-sigmoidal curve fitting. PeerJ. doi: 10.7717/peerj.4251

Deolu-Ajayi AO, Meyer AJ, Haring MA, Julkowska MM, Testerink C (2019) Genetic Loci Associated with Early Salt Stress Responses of Roots. iScience 21: 458–473

Duan L, Dietrich D, Ng CH, Chan PMY, Bhalerao R, Bennett MJ, Dinneny JR (2013) Endodermal ABA signaling promotes lateral root quiescence during salt stress in Arabidopsis seedlings. Plant Cell 25: 324–41

Geng Y, Wu R, Wee CW, Xie F, Wei X, Mei P, Chan Y, Tham C, Duan L, Dinneny J (2013) A Spatio-Temporal Understanding of Growth Regulation during the Salt Stress Response in Arabidopsis. Plant Cell 2132–2154

Gigli-Bisceglia N, van Zelm E, Huo W, Lamers J, Testerink C (2022) Arabidopsis root responses to salinity depend on pectin modification and cell wall sensing. Development 149: 1–13

Hassani A, Azapagic A, Shokri N (2020) Predicting long-term dynamics of soil salinity and sodicity on a global scale. Proc Natl Acad Sci U S A 117: 33017–33027

Hassani A, Azapagic A, Shokri N (2021) Global predictions of primary soil salinization under changing climate in the 21st century. Nat Commun 12: 1–17

Julkowska MM, Hoefsloot HCJ, Mol S, Feron R, de Boer G-J, Haring M a, Testerink C (2014) Capturing Arabidopsis root architecture dynamics with ROOT-FIT reveals diversity in responses to salinity. Plant Physiol 166: 1387–1402

Julkowska MM, Koevoets IT, Mol S, Hoefsloot H, Feron R, Tester MA, Keurentjes JJB, Korte A, Haring MA, De Boer GJ, et al (2017) Genetic components of root architecture remodeling in response to salt stress. Plant Cell 29: 3198–3213

Julkowska MM, Testerink C (2015) Tuning plant signaling and growth to survive salt. Trends Plant Sci 20: 586– 594

Li H, Duijts K, Pasini C, Santen JE van, Wang N, Zeeman SC, Santelia D, Zhang Y, Testerink C (2022) Effective root responses to salinity stress include maintained cell expansion and carbon allocation. bioRxiv 2022.09.01.506200

Lobet G, Pagès L, Draye X (2011) A novel image-analysis toolbox enabling quantitative analysis of root system architecture. Plant Physiol 157: 29–39

Lucas M, Godin C, Jay-Allemand C, Laplaze L (2008) Auxin fluxes in the root apex co-regulate gravitropism and lateral root initiation. J Exp Bot 59: 55–66

Munns R, Tester M (2008) Mechanisms of salinity tolerance. Annu Rev Plant Biol 59: 651–81

Péret B, Li G, Zhao J, Band LR, Voß U, Postaire O, Luu D, Ines O Da, Casimiro I, Lucas M, et al (2012) Auxin regulates aquaporin function to facilitate lateral root emergence. Nat Cell Biol 14: 991–998

Rus A, Baxter I, Muthukumar B, Gustin J, Lahner B, Yakubova E, Salt DE (2006) Natural variants of AtHKT1 enhance Na+ accumulation in two wild populations of Arabidopsis. PLoS Genet 2: 1964–1973

Voß U, Wilson MH, Kenobi K, Gould PD, Robertson FC, Peer WA, Lucas M, Swarup K, Casimiro I, Holman TJ, et al (2015) The circadian clock rephases during lateral root organ initiation in Arabidopsis thaliana. Nat Commun. doi: 10.1038/ncomms8641

Waidmann S, Sarkel E, Kleine-Vehn J (2020) Same same, but different: Growth responses of primary and lateral roots. J Exp Bot 71: 2397–2411

West G, Inze D, Beemster GTS (2004) Cell Cycle Modulation in the Response of the Primary Root of Arabidopsis to Salt Stress. Plant Physiol 135: 1050–1058

Van Zelm E, Zhang Y, Testerink C (2020) Salt Tolerance Mechanisms of Plants. Annu Rev Plant Biol 71: 403–433

Zou Y, Gigli-Bisceglia N, van Zelm E, Julkowska MM, Zeng Y, Cheng Y, Koevoets IT, Jørgensen B, Giesbers M, Vroom J, et al (2022a) Cell wall extensin arabinosylation is required for root directional response to salinity. bioRxiv 497042

Zou Y, Zhang Y, Testerink C (2022b) Root dynamic growth strategies in response to salinity. Plant Cell Environ 45: 695–704

